# proABC-2: PRediction Of AntiBody Contacts v2 and its application to information-driven docking

**DOI:** 10.1101/2020.03.18.967828

**Authors:** F. Ambrosetti, T. H. Olsen, P. P. Olimpieri, B. Jiménez-García, E. Milanetti, P. Marcatilli, A.M.J.J. Bonvin

**Author notes:** authors contributed equally (F. Ambrosetti, T. H. Olsen).

## Abstract

Monoclonal antibodies (mAbs) are essential tools in the contemporary therapeutic armoury. Understanding how these recognize their antigen is a fundamental step in their rational design and engineering. The rising amount of publicly available data is catalysing the development of computational approaches able to offer valuable, faster and cheaper alternatives to classical experimental methodologies used for the study of antibody-antigen complexes.

Here we present proABC-2, an update of the original random-forest antibody paratope predictor, based on a convolutional neural network algorithm. We also demonstrate how the predictions can be fruitfully used to drive the docking in HADDOCK.

The proABC-2 server is freely available at: https://bianca.science.uu.nl/proabc2/.

## 1. Introduction

Monoclonal antibodies (mAbs) are now well established in the contemporary therapeutic repertoire. Indeed, in 2018, 12 antibodies were granted first approval by either the European Medicines Agency (EMA) or by the Food and Drug Administration (FDA) while about 570 are undergoing clinical development at various stages (Kaplon and Reichert, 2019). The reasons behind the increasingly consolidated use of mAbs as therapeutics should be sought in their high affinity and specificity toward their cognate antigen and their modular architecture which facilitates their engineering (Chames *et al*., 2009). Understanding the fundamentals of antibody-antigen interactions is a critical step for the rational design and engineering of immunoglobulins. Since classical experimental approaches used to characterize antibodies (e.g. NMR, x-ray, mass spectrometry) are often expensive and time consuming, computational tools offer valuable and complementary approaches which can provide information at different levels (sequence and/or structural) (Norman *et al*., 2019).

To this end, we previously reported a method named proABC (Olimpieri *et al*., 2013) that can predict antibody residues forming intermolecular contacts with the cognate antigen, as well as the nature of their contacts, distinguishing between hydrogen bonds and hydrophobic interactions. proABC is based on a random forest algorithm, using the antibody heavy and light chain sequences, the hypervariable loop canonical structures and lengths (Chothia and Lesk, 1987) and the germline family as features (Schatz and Swanson, 2011). Its performance has been validated by us (Olimpieri *et al*., 2013) and others (Peng *et al*., 2014) demonstrating good accuracy and reliability.

Here we present proABC-2, an update of the original algorithm, using the same set of features, but based on a deep learning framework shown to be successful in achieving similar goals (Liberis *et al*., 2018; Deac *et al*., 2019). Furthermore, we show how the proABC-2 predictions can be used to drive the modelling of antibody-antigen complexes using the information-driven docking approach HADDOCK (Van Zundert *et al*., 2016), which was recently demonstrated to be the best option of the compared methods for antibody-antigen modelling (Ambrosetti *et al*., 2020). proABC-2 is integrated in a freely available web server that predicts paratope residues forming general contacts as well as those involved in hydrogen bonds and hydrophobic interactions.

## 2. Methods

### 2.1. Dataset

The full protein data bank (PDB) was scanned using in-house Hidden Markov Models (HMM) in order to identify all the antibody structures deposited. Immunoglobulins having only one chain (nanobodies), a resolution higher than 3Å or not solved with an antigen were excluded. Finally, using *cd-hit* (Fu *et al*., 2012) all the antibodies sharing a sequence identity higher than 95% with any other immunoglobulin of the dataset were removed.

We ended up with a dataset of 769 complexes (*CNN-dataset*) which was used to train the model. For the docking studies a dataset of 16 complexes, all with available unbound structures, corresponding to the new antibody-antigen entries of the protein-protein benchmark version 5.0 was used (Vreven *et al*., 2015).

Moreover, for a fair comparison with Parapred, the same structures used to train it (*Parapred-set*) have been used to train proABC-2.

### 2.2. Interaction calculation

For all the complexes of the CNN-dataset, the non-covalent interactions including intermolecular hydrogen bonds and hydrophobic interactions were calculated. Non-covalent interactions were determined using a distance cut-off of 3.9Å. Hydrogen bonds were calculated by defining donors (D) as any N/O/F/S connected to a hydrogen atom and acceptors (A) as any N/O/F/S within a distance threshold (2.5Å) of that hydrogen and by filtering the matches for D-H-A triplets with a minimum angle of 120 degrees (Baker and Hubbard, 1984). Finally, hydrophobic interactions were computed using a distance cut-off of 4.4Å between any heavy atom of two hydrophobic residues (Bissantz *et al*., 2010).

General contacts were calculated using an in-house R script while H-bond and hydrophobic interactions were determined using *interfacea* (Rodrigues *et al*., 2019).

### 2.3. Neural network features

In order to train the CNN a specific set of features was used:

1. *Light and heavy chain sequences* aligned using HMM profiles. In particular for the H3 alignment insertions were positioned in the middle between the two conserved residues Cys92 and Gly104 according to the previously described method (Morea *et al*., 1998). Each sequence position was considered as a variable. To allow the textual information of a sequence to be processed by an algorithm, each residue was converted into numerical values using one-hot encoding, the representation of categorical variables (i.e. a residue) as binary vectors. Here, a 20×1 vector has been used consisting of all zeros except at the index of the given residue, which was marked with a 1. Concurrent, a 20×1 vector of only zeros represented a gap. The heavy and light chains were represented by a 297×20 array.
2. *Hypervariable loops canonical structures* calculated according to the key residues found within and outside the loops (Chothia and Lesk, 1987; Vargas-Madrazo and Paz-García, 2002; Morea *et al*., 1998). One-hot encoding was used.
3. *Length of the hypervariable loops* defined according to the Chothia numbering scheme.
4. *Germline family* and *source organism* determined using *igblastp* (Ye *et al*., 2013). One-hot encoding was used.

### 2.4. Convolutional Neural Network (CNN)

The neural network used by proABC-2 consists of three convolutional modules (Conv11, Conv12 and Conv2), a fully connected feed-forward module (Ff1) and an output layer (Figure S1). Conv11 and Conv12 are identical and consist of three parts; a 1D convolutional layer with 32 filters of size 3×1 and a stride of 1, followed by a 1D max pooling layer of size 10×1 and a stride of 3 and finally a dropout layer with a dropout rate of 0.15. Conv2 also consists of three parts; a 1D convolutional layer with 64 filters of size 3×1 and a stride of 1, followed by a 1D average pooling layer of size 6×1 and a stride of 3 and finally a dropout layer with a dropout rate of 0.15. Ff1 consists of a fully connected layer with 512 nodes followed by a dropout layer with a dropout rate of 0.10. The final output layer has for each of the 297 residues 3 nodes, predicting the general interactions, H-bonds and hydrophobic interactions, amounting to 891 nodes. The model was constructed using the python package *Tensorflow* (Abadi *et al*., 2016).

These modules are combined in the following way. The one-hot encoded heavy and light chains are connected to Conv11 and Conv12 respectively. The extracted features of the heavy and light chains are then concatenated and enter Conv2 for a deeper feature extraction. The final extracted features from Conv2 are then flattened (reduced to one dimension) and concatenated with the additional features (germline, loop lengths and canonical structures) before entering Ff1 and finally into the output nodes. The purpose of Ff1 is to learn each individual residue’s role in the paratope based on the extracted features and the additional ones. The architecture is shown in Figure S1. The network was optimized with a focal loss (Lin *et al*., 2017) and a stochastic gradient descent (SGD) optimizer. The learning rate followed a one-cycle learning rate policy (Smith and Topin, 2017) with a max learning rate of 0.5, a minimum learning rate of 0.1% of the max one and maximum momentum of 0.9. Exponential Linear Units (ELU) were used as activation functions for Conv11, Conv12, Conv2 and Ff1, and sigmoid on the final output. Dropout (Srivastava *et al*., 2014) and early stopping (Prechelt, 1998) were used throughout training as regularization techniques. All hyperparameters (i.e. nodes, filter sizes, dropout rate etc.) mentioned above were found empirically.

### 2.5. Model evaluation

The evaluation of the model has been performed using 10-fold nested cross validation on the full CNN-set (769 complexes). The performance was measured taking into account three different metrics: area under the Receiver Operating Characteristic curve (AUC), Matthew Correlation Coefficient (MCC) and F-score. MCC and F-score were calculated using a threshold of 0.40, 0.30 and 0.30, respectively for Pt, Hy and Hb.

These cut-offs were selected by averaging the thresholds that for each fold of the cross validation gave the best MCC.

### 2.6. Docking scenarios and settings

To assess the impact of the proABC-2 predictions on HADDOCK’s docking performance the following docking scenarios were evaluated:

1. *Pred Para – Surf*: No previous information about the epitope is provided to HADDOCK. The docking was performed by using the residues predicted to be in contact by proABC-2, defined as active, and the antigen residues having a relative accessible surface areas (RSA) ≥ 40% calculated with NACCESS (Hubbard SJ, 1993), provided as passive. Default docking settings were used except for the sampling that was increased to 10000, 400, 400 for it0, it1 and water respectively.
2. *Pred Para – Epi 9*: In this case we use a loose definition of the epitope region by selecting all the antigen residues within a 9Å distance from the antibody in the reference structure. We provide to HADDOCK the residues predicted by proABC-2 as active and the defined antigen residues as passive. Default docking settings were used except for the sampling that was increased to 5000, 400, 400 for it0, it1 and water respectively.

The antibody structures were renumbered with an in-house R script using a consecutive numbering as HADDOCK is not able to deal with the insertion format of the Chothia scheme.

### 2.7. Docking evaluation criteria

Docking performance was evaluated by classifying the models into 3 classes: high (***), medium (**) of low (*) quality defined according to the CAPRI criteria (Janin et al., 2003; Méndez et al., 2003) (see Table S2). We calculated the interface root mean square deviation (i-RMSD), the ligand root mean square deviation (L-RMSD) and the fraction of native contacts (F_nat_) as already reported (Méndez *et al*., 2003). Briefly, the i-RMSD is calculated on the interface residues backbone atoms defined on the native structure using a 10Å cut-off, the L-RMSD is calculated by superimposing on the antibody backbone atoms and calculating the RMSD on the antigen ones. Finally, F_nat_ is calculated as the number of native contacts in a docking model divided by the total number of contacts in the reference structure. These are defined using a 5Å cut-off.

F_nat_ has been calculated using in-house scripts while fitting and RMSD calculations were performed using the McLachlan algorithm (McLachlan, 1982) as implemented in the program ProFit (http://www.bioinf.org.uk/software/profit/) from the SBGrid distribution (Morin *et al*., 2013).

## 3. Results

### 3.1. proABC-2 performance

The prediction performance of proABC-2 was measured, after a 10-fold nested cross-validation, in terms of AUC, MCC and F-score values for all the general interactions of the paratope (*Pt)*, hydrophobic interactions (*Hy*) and for hydrogen bonds (*Hb*). The highest performance is obtained for *Pt* (0.96, 0.57 and 0.59 respectively for AUC, MCC and F-score) and decreases for *Hy* (0.95, 0.44 and 0.41) and *Hb* (0.94, 0.33 and 0.27). This is due to the smaller number of *Hb* and *Hy* interactions in the training set compared to the general (*Pt*) ones.

### 3.2. Comparison with Parapred

For a fair comparison proABC-2 was trained on the *Parapred-set* and the AUC, MCC and F-score were calculated on the same residues used by Parapred to make the predictions (CDRs defined according to the Chothia numbering scheme plus two extra residues at both ends) after a 10-fold nested cross validation. For proABC-2 the MCC and F-score were calculated using a threshold of 0.37 (determined as explained in paragraph 2.5 of the Method section), while the values from the work of Liberis et al. (Liberis *et al*., 2018) are reported for Parapred. The results in Table 1 show that proABC-2 outperforms Parapred in terms of AUC and MCC but has a lower performance in terms of F-score.

**Table 1:** Classification of docking models in the classes: ***, **, * according to F_nat_, and either L-RMSD or i-RMSD measures.

| Class | $F_{\text{nat}}$ | L-RMSD[Å] | i-RMSD[Å] |
| --- | --- | --- | --- |
| High (***) | □ 0.5 | ≤ 1.0 | or ≤ 1.0 |
| Medium (**) | □ 0.3 | ≤ 5.0 | or ≤ 2.0 |
| Acceptable (*) | □ 0.1 | ≤ 10.0 | or ≤ 4.0 |

**Table 2:** Performance comparison between Parapred and proABC-2.

| Method | AUC | MCC | F-score |
| --- | --- | --- | --- |
| *proABC-2* | 0.91 | 0.56 | 0.62 |
| *Parapred* | 0.88 | 0.55 | 0.69 |

### 3.3. Prediction-driven docking accuracy

We investigated whether the predictions obtained from proABC-2 can be used to drive antibody-antigen docking using the HADDOCK 2.2 webserver (Van Zundert *et al*., 2016). For unbiased predictions, the model was trained excluding all sequences sharing ≥95% sequence identity with any structure used for docking. Only residues predicted as *Pt* were used for docking (using a 0.40 cutoff). The results were compared to a previous study performed using the hypervariable loops (Ambrosetti *et al*., 2020)(see Figures 2 and 3). The performance was evaluated in terms of success rate defined as the number of complexes for which at least one acceptable, medium or high-quality complex was found in the top 1, 5, 10, 20, 50 and 100 ranked models. Figure 2 shows the results of the docking obtained by providing to the algorithm all solvent accessible residues of the antigen and either the antibody hypervariable loops (HV-Surf) or the proABC-2 predictions (*Pt*) (Pred-Surf). The HV-Surf docking led to slightly better results for the top 1,5 and 10 with 25.0%, 31.2% and 31.2% success rates respectively, compared to 18.7%, 25.0% and 25.0% for Pred-Surf. The proABC-2 predictions give better results for the top 50 and 100 (50% and 62.5% success rates, respectively). Thus, even if HADDOCK is able to generate correct models, the scoring is not able to rank them in the top. As for the quality of the docking models, using the HV loop leads to better quality models overall.

**Figure 1:**
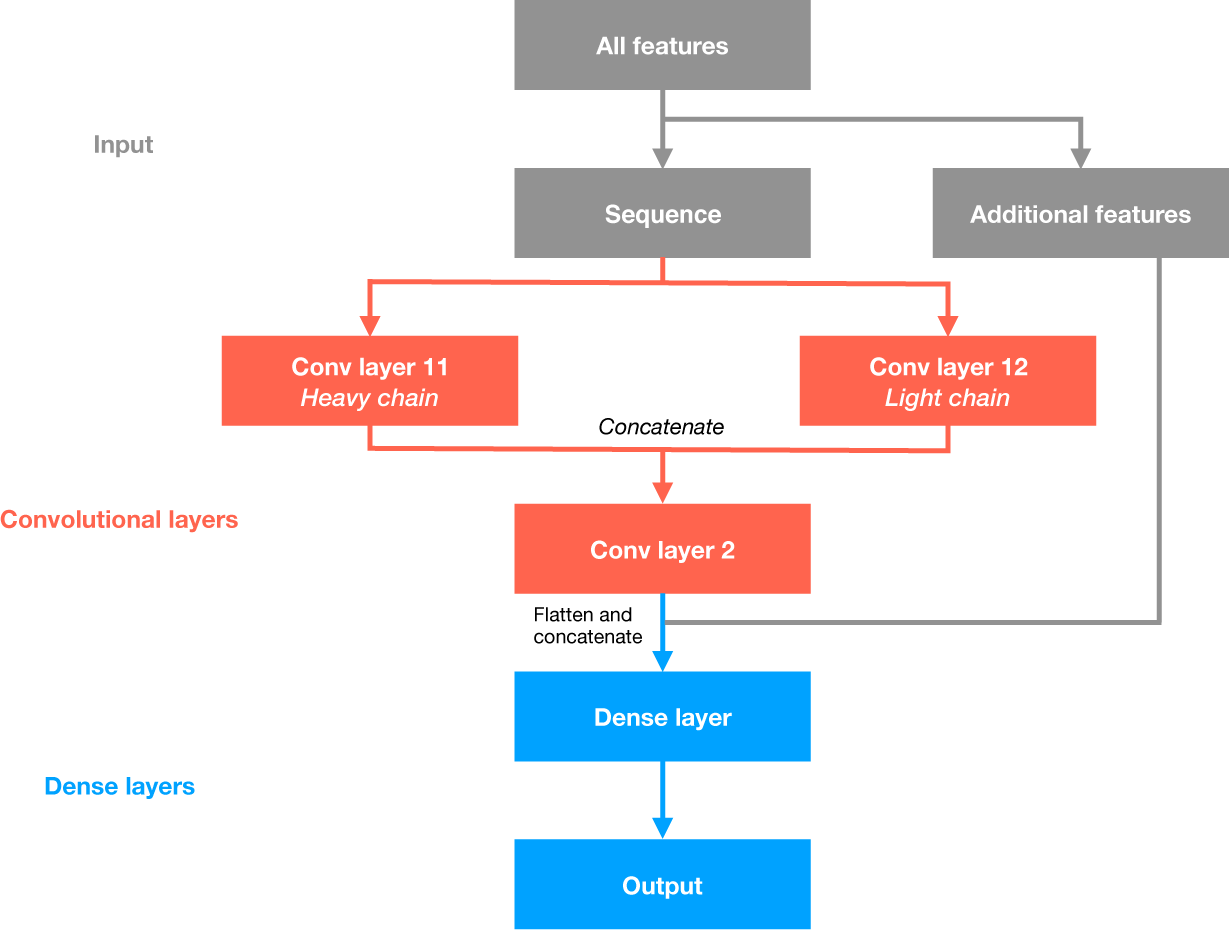
The CNN architecture implemented in the proABC-2 method.

**Figure 2:**
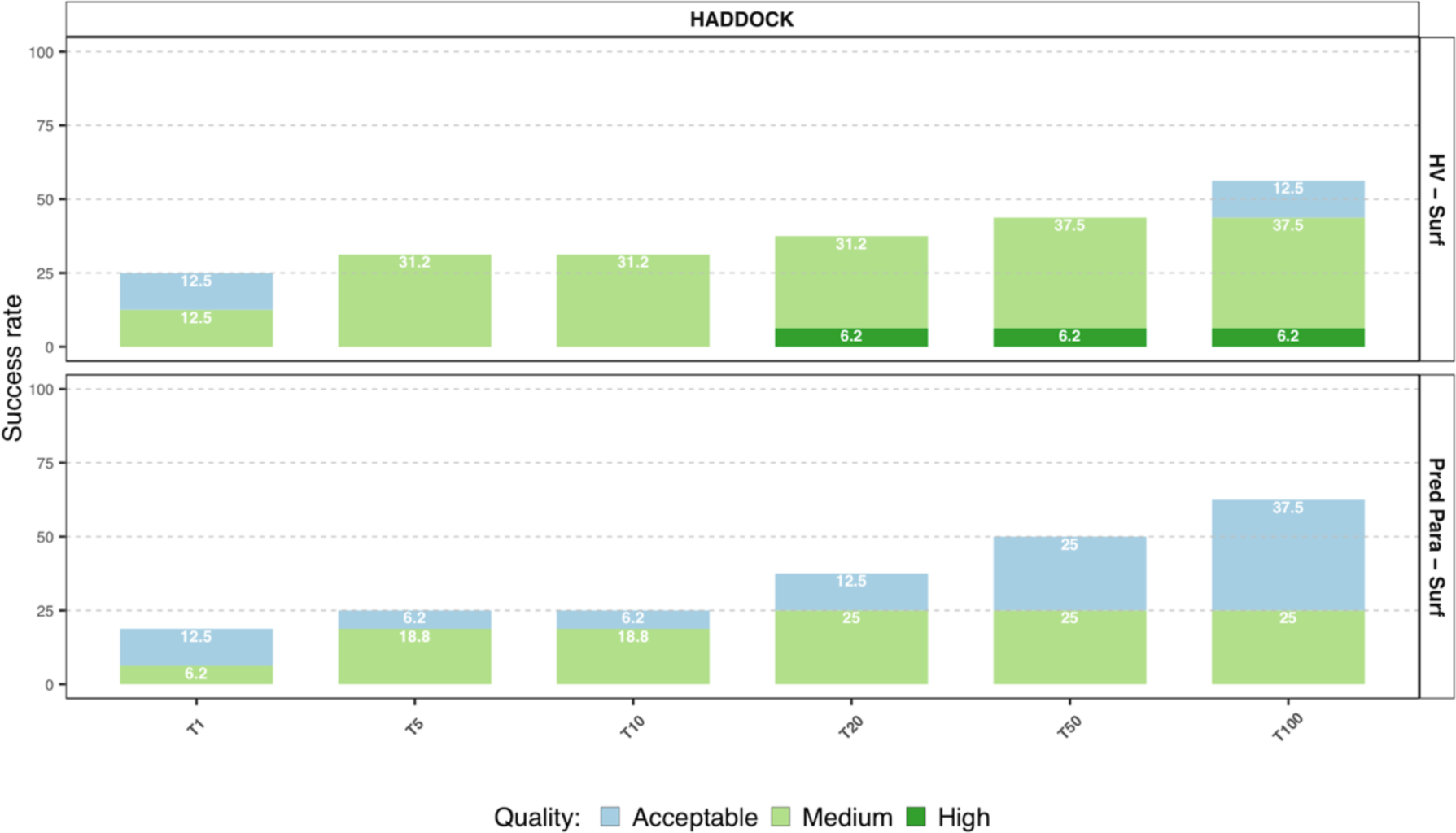
HADDOCK success rate as a function of the top 1, 5, 10, 20, 50 and 100 ranked models. The top row (HV - Surf) shows the success rate using the antibody HV loops and the entire antigen surface as restraints. The second represents the success rate achieved by driving the docking with the proABC-2 predictions and the full surface of the antigen. The colour coding indicates the quality of the models according to CAPRI criteria.

**Figure 3:**
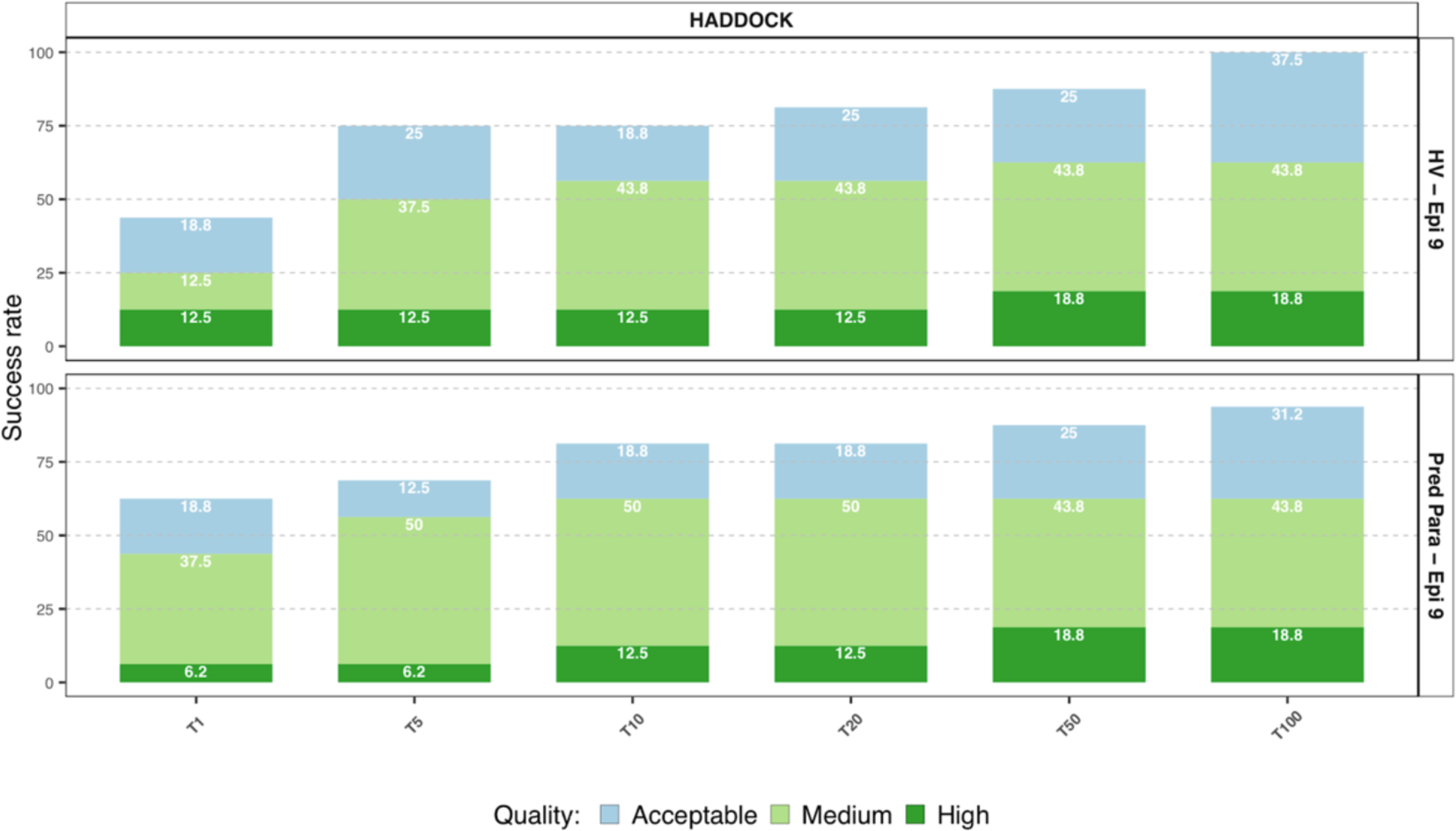
HADDOCK success rate as a function of the top 1, 5, 10, 20, 50 and 100 ranked models. The top row (HV – Epi 9) shows the success rate using the antibody HV loops and a loose definition of the epitope using a 9Å cut-off. The second represents the success rate achieved by driving the docking with the proABC-2 predictions and the same definition for the epitope on the antigen. The colour coding indicates the quality of the models according to CAPRI criteria.

Figure 3 shows the results of the docking obtained by providing to the algorithm a loose definition of the epitope following the definition given in (Ambrosetti *et al*., 2020). In this scenario, the proABC-2 predictions led to a remarkable improvement of the Top1 success rate from 43.8% (using HV) to 62.5%. In general, the use of the proABC-2 predictions resulted in an improvement of the quality of the generated models, mainly reflected in the number of medium quality ones.

### 3.4. Web server

proABC-2 is freely available as a web server at https://bianca.science.uu.nl/proabc2. It only requires the sequences of the heavy and light chains. The input is processed to calculate all of the sequence-derived features (germline, canonical structures and length of the HV loops) and these are passed to the CNN to make the predictions. The computation only takes a few seconds. The results page reports in a bar plot the residue probabilities of making a general, H-bond and hydrophobic interactions (see Figure 4). Two files (for the heavy and light chains) are provided as output, containing for each residue the different probabilities.

**Figure 4:**
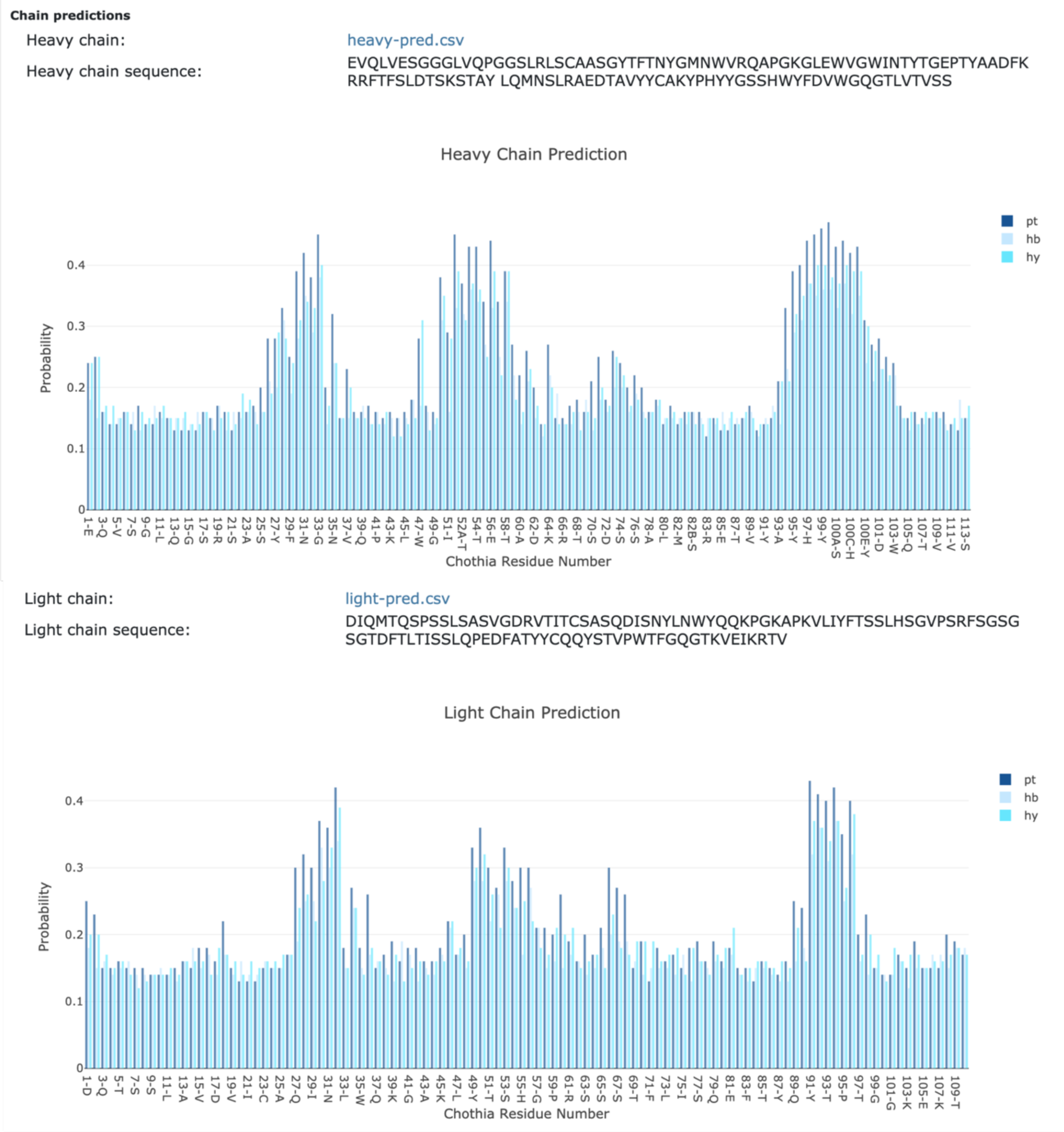
Output page of the proABC-2 web server (https://wenmr.science.uu.nl/proabc2/). It shows the interaction probability of the antibody residues belonging to the heavy and light chain.

## 4. Conclusions

proABC-2 is based on a deep learning framework and shows a high performance with an AUC of 0.96 and an MCC of 0.57. Its predictions should be useful for antibody design such as in silico affinity maturation or humanization. We also demonstrated how these predictions can guide molecular docking, showing in particular that if a loose definition of the epitope region is provided, the proABC-2 predictions leads to improvements of both success rate and quality of the docked models. This suggests that different strategies might be followed depending on the available information about the epitope. To our knowledge proABC-2 is the only available method, specifically designed for antibodies, able to predict the paratope residues along with the type of interaction. The method is freely available as a web server and provides a straightforward user-friendly interface.

## Funding

This work has been supported by the European Union Horizon 2020 BioExcel (project 823830).

## Conflict of Interest

none declared.

